# Antifungal activity of *Cinnamomum cassia* and *Origanum compactum* essential oils against *Aspergillus oryzae*

**DOI:** 10.1101/2022.03.24.485625

**Authors:** Manon Leclercq, Isciane Commenge, Marylou Bouriot, Floricia Crusset, Patrick Gonzalez, Julien Grimaud, Frank Yates, Jacqueline Sarfati-Bert, Agnès Saint-Pol

## Abstract

Aspergillosis is a nosocomial disease that usually affects the respiratory system. While most patients develop mild symptoms, aspergillosis can become a serious health threat in immunocompromised populations and patients with pre-existing respiratory conditions. Aspergillosis is caused by fungi of the *Aspergillus* genus. Existing treatments include drugs such as amphotericin B, with strong side effects. Numerous studies have shown the antifungal properties of essential oils (EOs), which represent potential alternative treatments against fungal infections. Here, we screened the antifungal properties of five EOs against *Aspergillus oryzae*: *Melaleuca alternifolia, Mentha x piperita, Thymus zygis, Origanum compactum*, and *Cinnamomum cassia*. Of the five EOs, two demonstrated antifungal activity: *Origanum compactum* acted as a fungistatic, while *Cinnamomum cassia* showed both fungistatic and fungicidal effects against *A. oryzae*. Therefore, both EOs represent potential alternative treatments against Aspergillosis.

## Introduction

Aspergillosis is a nosocomial infection of the respiratory tract. It is caused by members of the genus *Aspergillus. Aspergillus* conidia are found everywhere in the environment (air, soil, plants, water…) as well as inside houses and hospitals, in which they find suitable conditions for their development. Patients get infected by inhaling fungi which then develop inside the lungs [1,2].

Depending on the host’s immune system, aspergillosis may cause various symptoms. For instance, in immunocompromised patients, it usually leads to a lung disease named invasive pulmonary aspergillosis, which is the most common cause of death by aspergillosis. Other aspergillosis related pathologies exist, such as allergic bronchopulmonary aspergillosis (ABPA), which occurs in atopic patients [2,3].

While many species of the genus *Aspergillus* may cause aspergillosis, one of the most reported pathogens is *Aspergillus fumigatus* [1,2]. *A. fumigatus* is a Class 2 pathogenic agent [4]. Therefore, scientists who wish to study it must gain access to research facilities of an equivalent biosafety level.

While *A. fumigatus* is the main cause of aspergillosis, other fungi of the genus *Aspergillus* can also cause it. For example, *Aspergillus oryzae* is often reported in cases of ABPA [5,6]. *A. oryzae* is classified as a Class 1 pathogenic agent [7], which means it may be handled in any research facilities with minimal biosafety level.

Furthermore, various *Aspergillus* genomes share a high degree of similarity, suggesting that this species is phylogenetically very close to *A. fumigatus* [8]. The percentage of similarity between *A. fumigatus* and other *Aspergillus* species such as *A. flavus, A. niger, A. oryzae* and *A. nidulans* has been studied at the amino acid level, where researchers found 78% of similarity between *A. fumigatus* and *A. oryzae* - higher than the similarity between *A. fumigatus* and other the *Aspergillus* species reported [9] (**Table 1**). It is worth noting that some studies suggest that both *A. fumigatus* and *A. oryzae* may be able to reproduce asexually [10], although this last point is subject to debate [11]. Therefore, *A. oryzae* represents a good alternative model to *A. fumigatus*, for scientists who wish to study aspergillosis agents with no access to high biosafety level facilities.

**Table 1:** Percentage of similarity from an alignment of 2753 orthologous proteins across *Aspergillus* genomes. *A. oryzae* and *A. fumigatus* show the highest percentage of alignment. Adapted from [9].

|  | *A. flavus* | *A. niger* | *A. oryzae* | *A. nidulans* |
| --- | --- | --- | --- | --- |
| *A. fumigatus* | 77.5% | 76.6% | 78% | 73.9% |

Nowadays, aspergillosis is treated with antifungal drugs, which usually consist of ergosterol interfering agents (Amphotericin B^®^) and / or echinocandins (Caspofungin^®^) [2]. However, these drugs have serious side effects such as renal insufficiency and allergic reactions [5]. In addition, *in vitro* approaches have shown that *A. fumigatus* might still grow, even in the presence of these drugs at high concentrations [2,12]. Therefore, there is a need for alternative treatments to aspergillosis that would be both less toxic for the patients and more effective.

Our goal was to identify new candidate drugs against aspergillosis. Recently, various studies have revealed the antifungal properties of essential oils (EOs) [3,13–18]. Therefore, we decided to test five EOs against *A. oryzae*: *Mentha x piperita, Melaleuca alternifolia, Thymus zygis, Origanum compactum, Cinnamomum cassia*.

Most of these EOs have known activity against Aspergillus strains. For instance, *Mentha x piperita* appeared effective against *A. flavus* and *A. parasiticus* [14]. *Melaleuca alternifolia* showed antifungal activity against *A. niger* [15]. *Thymus zygis* and *Origanum compactum* revealed antifungal properties against both *A. flavus* and *A. niger* [16]. *Oregano Compactum* contains a phenolic compound called carvacrol which could explain its inhibiting activity on some *Aspergillus* strains [17]. Finally, *Cinnamomum cassia* also showed a high antifungal activity against *A. niger*, probably because it contains cinnamaldehyde, an antifungal molecule [18].

Here we investigated both the fungicidal and fungistatic activity of these five EOs at various concentrations against A. oryzae. *Melaleuca alternifolia, Mentha x piperita and Thymus zygis* showed antifungal activity but only at high concentrations (over 1mg/mL), after 48h of incubation with EOs. *Origanum compactum* proved an interesting candidate, inhibiting the growth of *A. oryzae* at a lower concentration (700μg/mL) thanks to fungistatic activity. The EO that proved most effective was *Cinnamomum cassia* which inhibited *A. oryzae* growth at 50μg/mL by fungistatic and fungicidal action, its efficacy depending on the incubation time.

## Materials and Methods

### Fungal strain

*Aspergillus oryzae var. oryzae* CBS 816.2 was purchased from the CBS-KNAW Fungal Biodiversity Centre (Netherlands).

### Preculture

Prior to any experiment, we prepared fungal precultures as follows: *A. oryzae* was seeded into a Czapek Agar medium (0.01g/L ferrous sulfate, 0.5g/L magnesium sulfate, 0.5g/L potassium chloride, 1g/L potassium phosphate, 3g/L sodium nitrate, 30g/L D-sucrose, 12g/L agar) containing 1M of KCl. The preparation was incubated at 30°C for 6 days. The fungi were then collected with 0.05% Tween-20^®^ (ref. P1379, Sigma Aldrich, Burlington MA, USA). Finally, the fungal concentration was adjusted to 10^5^ conidia/mL.

### Antifungal screening

We prepared the following dilutions in Sabouraud medium (10g/L peptone, 20g/L glucose) containing 0.05% Tween-20^®^:

- *Melaleuca alternifolia* (EAN 5420008503917, Pranarôm, Ghislenghien, Belgium): 1mg/mL; 2mg/mL; 3mg/mL; 4mg/mL.
- *Mentha x piperita* (EAN 3401560104783, PurEssentiel, Bruxelles, Belgium): 1mg/mL; 2mg/mL; 3mg/mL; 4mg/mL.
- *Thymus zygis* (EAN 3401599455122, Pranarôm, Ghislenghien, Belgium): 0.5mg/mL; 1mg/mL; 2mg/mL; 3mg/mL.
- *Origanum compactum* (EAN 3401599454002, PurEssentiel, Bruxelles, Belgium): 500μg/mL; 600μg/mL; 700μg/mL; 1000μg/mL.
- *Cinnamomum cassia* (EAN 5420008506826, Pranarôm, Ghislenghien, Belgium): 5μg/mL; 20μg/mL; 30μg/mL; 50μg/mL; 200μg/mL; 500μg/mL.
- *Ocimum basilicum* (EAN 3701056802378, PurEssentiel, Bruxelles, Belgium): 1mg/mL.
- Amphotericin B^®^ (ref. Y0000005, Sigma Aldrich, Burlington MA, USA): 2μg/mL.

For each EO dilution, *A. oryzae* was added (final working concentration in each well: 10^4^ conidia/mL). The mix was then poured in the well of a 24-well plate (final volume: 1ml per well). As additional controls, wells containing only the Sabouraud medium, or Sabouraud medium with 0.05% Tween-20^®^ were also seeded with 10^4^ conidia/mL (final volume: 1ml per well).

The wells were then incubated for 48h at 30°C. The results were observed with an inverted microscope (ref. AE31E, Motic).

### Fungicidal / fungistatic test

We prepared solid Sabouraud media (10g/L of peptone, 20g/L glucose, 20g/L agar) containing EOs or antibiotics at the following final concentrations:

- *Origanum compactum*: 700μg/mL, 800μg/mL, 900μg/mL.
- *Cinnamomum cassia*: 20μg/mL, 30μg/mL, 50μg/mL.
- *Ocimum basilicum* : 1mg/mL.
- Amphotericin B^®^: 2μg/mL.

The media were poured into Petri dishes (diameter 55mm, Sarstedt). As additional controls, Petri dishes containing only the solid Sabouraud medium, or the solid Sabouraud medium with 0.05% Tween-20^®^ were also prepared.

A 6mm diameter cellulose disk (Merck) was then soaked in a preculture of 10^4^ conidia/mL of *A. oryzae* and laid at the center of each Petri dish. The dishes were incubated for either 48h or 72h. After this first incubation, each diffusion disk was moved onto the center of a new Petri dish containing fresh Sabouraud solid medium and incubated for 48h.

The diameter of the growth disk was measured using a ruler at various times: end of first incubation, 24h after the start of the second incubation, and end of second incubation. All measurements of 6mm or less were considered as an absence of fungal growth, as 6mm was the diameter of the cellulose disk alone.

## Results

### Screening of several EOs

To determine which EOs had an antifungal activity, we incubated *A. oryzae* spores in the presence of the EOs of interest at various concentrations for 48h (**Figure 1**). *A. oryzae* germinates and forms a hyphal network in only 24h [19]. Therefore, for a given EO, if the growth medium is free of hyphae after 48h, then we can conclude that the EO is a potential inhibitor of *A. oryzae* development. On the contrary, if, after 48h, germination began or a network started to form, then we can consider that the EO has no observable antifungal activity against our strain of interest (**Figure 1A**).

**Figure 1:**
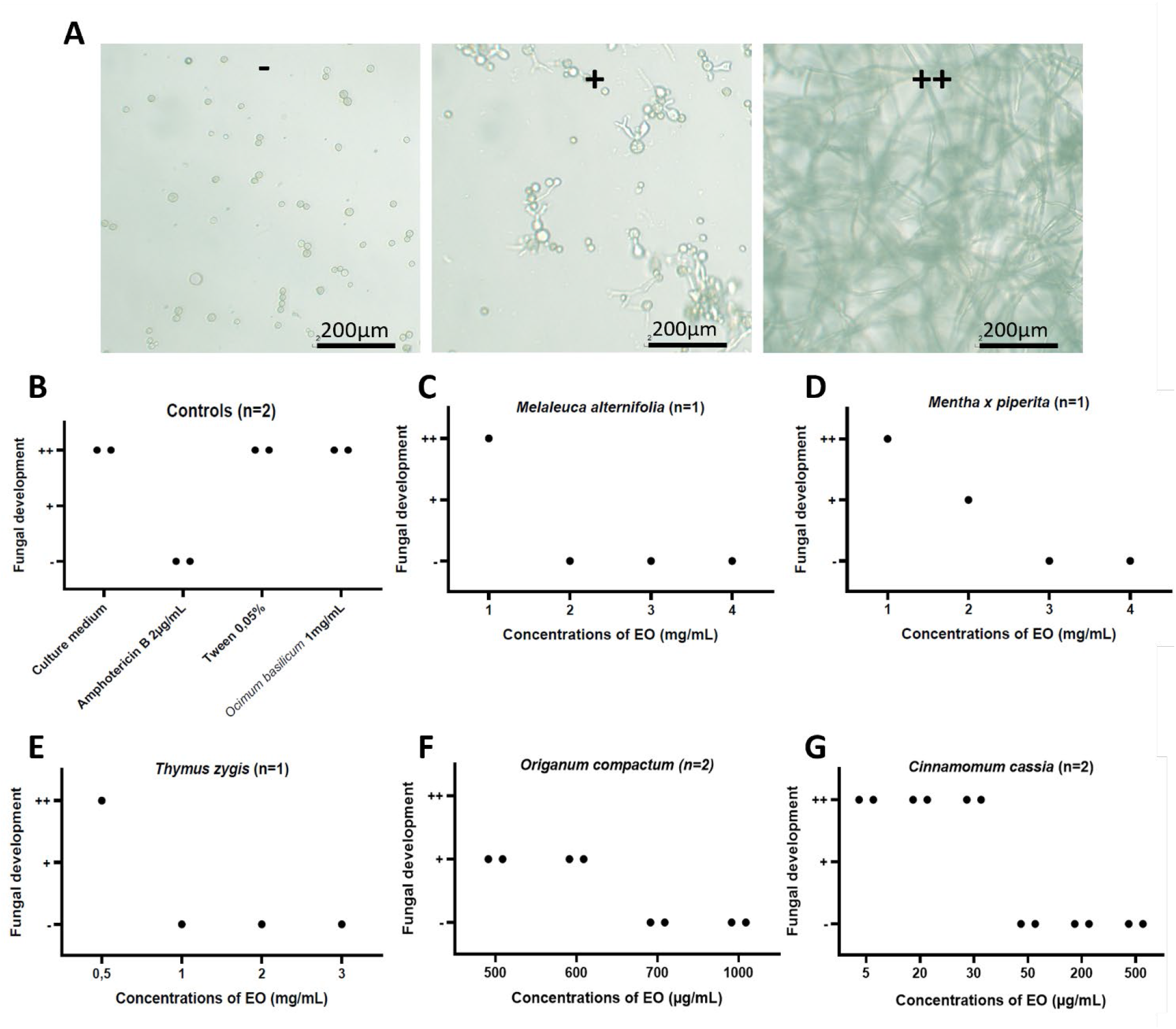
Screening of candidate EOs inhibiting *A. oryzae* growth. **(A)** Discrimination criteria of *A. oryzae* growth after 48h incubation with EOs or control treatments. For all panels, the fungal development will be scored as follows: (-) no development, (+) start of germination, (++) network formation. **(B)** Control conditions (culture medium alone; Amphotericin B; *Ocimum basilicum*; Tween-20), n=2 replicates per condition. **(C), (D)** Screening of *Melaleuca alternifolia*, and *Mentha x piperita* from 1mg/mL to 4mg/mL, n=1 replicate per condition. **(E)** Screening of *Thymus zygis* from 0,5mg/mL to 4mg/mL, n=1 replicate per condition. **(F)** Screening of *Origanum compactum* from 200μg/mL to 700μg/mL, n=2 replicates per condition. **(G)** Screening of *Cinnamomum cassia* from 5μg/mL to 500μg/mL, n=2 replicates per condition.

We carried out a set of controls (**Figure 1B**). First, to check the viability of our *A. oryzae* strain and the absence of potential antifungal activity of our growth medium alone, we performed the experiment in absence of EO. As expected, the fungus formed a network inside the wells. Second, we checked that we were able to prevent the development of *A. oryzae* with Amphotericin B, a known antifugal drug [20]. Finally, we tested an EO with no known antifungal activity: *Ocimum basilicum* [12]. As expected, this EO did not prevent fungal growth.

Then we screened our five EOs of interest (*Melaleuca alternifolia, Mentha x piperita, Thymus zygis, Origanum compactum*, and *Cinnamomum cassia*) at various concentrations, which were determined after preliminary tests (**Figure 1C-1G**).

Of these five EOs, three (*Melaleuca alternifolia, Mentha x piperita*, and *Thymus zygis*) showed antifungal activity only at concentrations higher than 1mg/mL (**Figure 1C-1E**). On the contrary, *Origanum compactum* and *Cinnamomum cassia* showed antifungal activity at concentrations as low as 700μg/mL for *Origanum compactum* and 50μg/mL for *Cinnamomum cassia* (**Figure 1F, 1G**).

Having identified two promising EO candidates with antifungal activity at relatively low concentrations (*Origanum compactum* and *Cinnamomum cassia*), we then decided to further investigate their antifungal properties.

### Analysis of fungistatic or fungicidal properties of selected EOs

We performed fungicidal and fungistatic tests with *Cinnamomum cassia* and *Origanum compactum* against *A. oryzae* (**Figure 2**). Briefly, we placed a small cellulose disk saturated with a solution of *A. oryzae* spores in a Petri dish containing a layer of solid Sabouraud medium mixed with the EO of interest (EO+). The Petri dish was incubated for 48 to 72h, after which we measured the diameter of fungal growth. Then, we transferred the same paper disk in a new Petri dish with fresh, EO-free medium (EO-) and measured, once again, the fungal growth after 24 and 48h of incubation (**Figure 2A, 2B**). Fungicidal substances would prevent fungal growth in both EO+ and EO-conditions, while fungistatic compounds would prevent fungal growth in the EO+ condition only.

**Figure 2:**
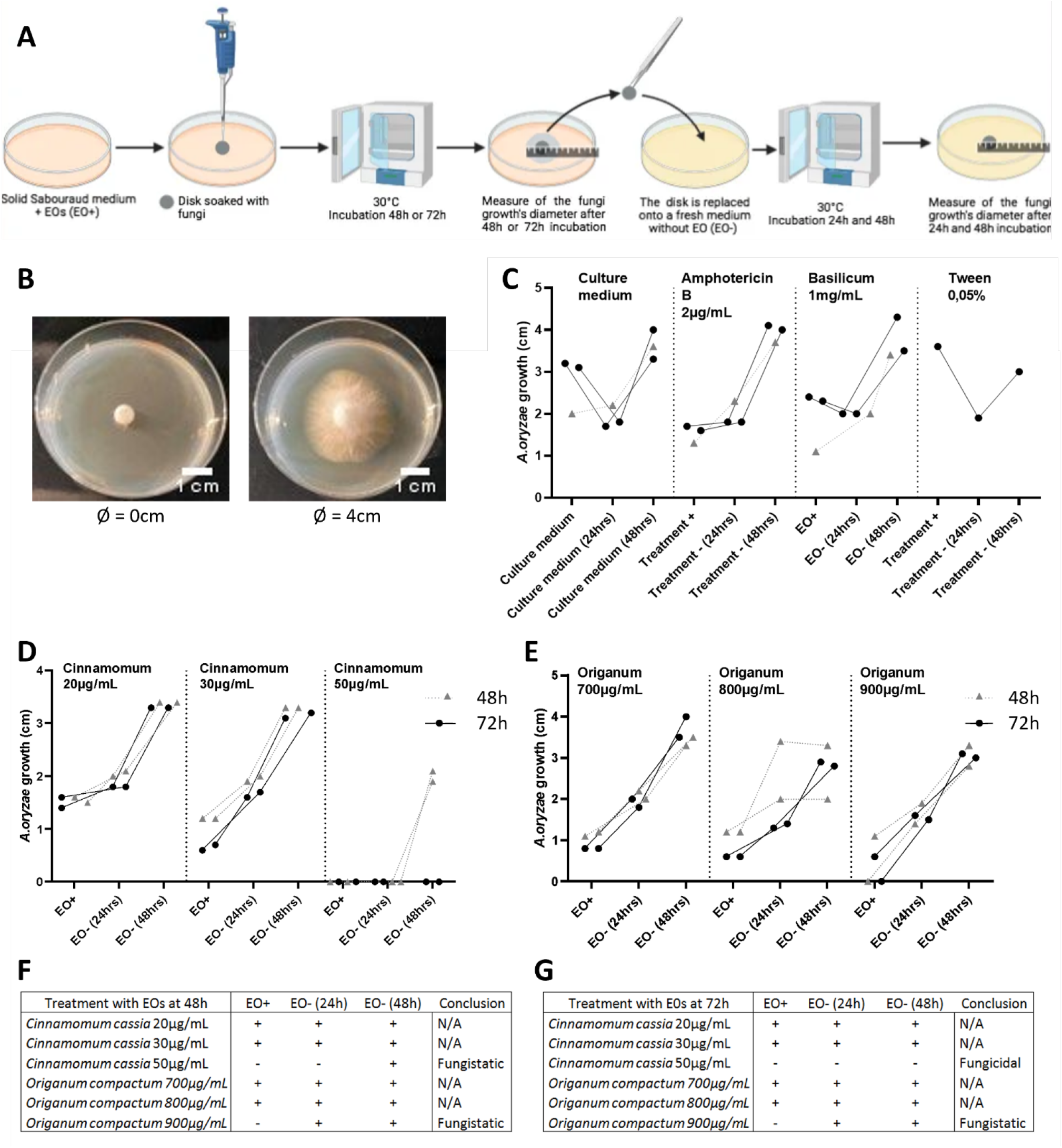
Fungicidal and fungistatic effects of EOs on *A. oryzae*. **(A)** Experimental approach. A cellulose disk saturated with A. oryzae is placed in a Petri dish containing solid Sabouraud medium with the EO of interest (EO+). The dish is incubated for 48 to 72h, after which fungal growth is measured. Then the disk is transferred in a new dish with EO-free medium (EO-). The fungal growth is measured at 24 and 48h of incubation. Fungicidal EOs would prevent growth in both EO+ and EO-conditions, while fungistatic EOs would prevent fungal growth in the EO+ condition only. **(B)** Exemplar growth disks. Left: no apparent growth. Right: 4cm growth. We measured fungal growth as the diameter of the disk after incubation. All diameters of 6mm or less were considered as an absence of fungal growth, as 6mm was the diameter of the cellulose disk alone. **(C)**. Control conditions (culture medium alone; Amphotericin B; *Ocimum basilicum*; Tween-20). For all graphs: all dots corresponding to the same cellulose disk are connected with a line. Grey triangles and dotted lines: the disks were left 48h in the EO+ medium. Black circles and solid lines: the EO+ incubation lasted 72h. **(D), (E)** Fungal growth in EO+ and EO-conditions. In (D), the EO+ medium contained *Cinnamomum cassia*, while it contained *Origanum compactum* in (E). **(F), (G)** Summary table of the measurements obtained from disks incubated for 48h (F) or 72h (G) in the EO+ condition. For all tables: (-) means an absence of fungal growth, while (+) represents an apparent growth.

To confirm our experimental approach, we performed a set of controls (**Figure 2C**). We examined the viability of our fungus strain and culture medium: as expected, A. oryzae formed a growth disk both before and after transfer to a new Petri dish. In addition, we confirmed the fungistatic effect of Amphotericin B [20], as well as the absence of antifungal activity of *Ocimum basilicum* [12] and Tween-20.

We then tested the two EOs with antifungal activity: *Cinnamomum cassia* and *Origanum compactum* (**Figure 2D-2G**). Both EOs inhibited the growth of *A. oryzae* in the EO+ condition at the highest concentration tested (*Cinnamomum cassia*: 50 μg/mL; *Origanum compactum*: 900 μg/mL), which confirmed the antifungal activity that we previously observed. Furthermore, the disks that were transferred from the EO+ to the EO-condition after 48h of incubation showed sizeable growth, indicative of a fungistatic effect from both *Cinnamomum cassia* and *Origanum compactum* (**Figure 2D-2F**).

However, after 72h of incubation in the EO+ condition, the two EOs showed different results. While we noted a fungistatic effect from *Origanum compactum*, the disk taken from *Cinnamomum cassia* EO+ medium developed no growth disk, which indicated a fungicidal effect (**Figure 2D, 2E, 2G**).

To conclude, we confirmed that both *Cinnamomum cassia* and *Origanum compactum* have antifungal properties against *A. oryzae*. We also showed the fungistatic effect of *Origanum compactum*, while *Cinnamomum cassia* acted both as a fungistatic and fungicidal agent, depending on the duration of the incubation in contact with the EO.

## Discussion

Aspergillosis is an infection of the respiratory tract, caused by members of the genus *Aspergillus* [1,2]. Current treatments have severe side effects and are not always effective [2,5,12]. Here, we explored EOs as alternative new treatments against aspergillosis. To do so, we screened the antifungal effect of five EOs against *A. oryzae*: *Mentha x piperita, Melaleuca alternifolia, Thymus zygis, Origanum compactum, Cinnamomum cassia*.

### Cinnamomum cassia, *a fungicidal EO against* A. oryzae

While all EOs showed at least some antifungal activity, not all of them were equally effective. Three EOs (*Melaleuca alternifolia, Mentha x piperita*, and *Thymus zygis*) prevented the growth of *A. oryzae* only at concentrations higher than 1mg/mL. On the contrary, *Origanum compactum* and *Cinnamomum cassia* showed antifungal activity at concentrations as low as 700μg/mL for *Origanum compactum* and 50μg/mL for *Cinnamomum cassia* (**Figure 1**).

All five EOs tested here are known for their antifungal activity against at least some members of the genus *Aspergillus* [14–18,21,22]. However, to our knowledge, no past study has assessed the effect of these EOs against *A. oryzae* specifically.

Furthermore, we found that both *Origanum compactum* and *Cinnamomum cassia* acted on *A. oryzae* as fungistatic agents. Interestingly, when the fungus was let in contact with the EO for 72h, *Cinnamomum cassia* also showed fungicidal properties (**Figure 2**).

The main antifungal ingredient of *Cinnamomum cassia* is cinnamaldehyde [18,21,22]. Other varieties of cinnamon based EOs exist, such as *Cinnamomum zeylanicum* (bark), which also contains cinnamaldehyde. A previous study found that *Cinnamomum cassia* was slightly better at inhibiting *A. niger* growth than *Cinnamomum zeylanicum* [18]. This is not surprising, considering the composition of these EOs: *Cinnamomum cassia* contains about 66% of cinnamaldehyde, while *Cinnamomum zeylanicum* contains 64% [18]. Therefore, one would expect *Cinnamomum cassia* to also show greater antifungal activity against *A. oryzae* than *Cinnamomum zeylanicum*. This hypothesis is open for testing.

### Cinnamomum cassia, *a non-toxic potential alternative to current aspergillosis treatment*

Today, aspergillosis is usually treated with one or two of the following drugs: Amphotericin B, an ergosterol interfering agent, and Caspofungin, an echinocandin [2]. However, these drugs have serious side effects such as nephrotoxicity and allergic reactions [5].

Furthermore, these drugs are not always effective, as *A. fumigatus* may still develop, even in the presence of these drugs [2,12]. Therefore, there is a need for new treatments that would bear fewer side effects and be more effective against aspergillosis.

Here we showed that *Cinnamomum cassia* is an effective fungicidal agent against *A. oryzae* at concentrations as low as 50μg/mL *in vitro*. Interestingly, it is nontoxic in clinical conditions [23]. Therefore, *Cinnamomum cassia* may represent a valuable alternative to conventional treatments against aspergillosis.

A recent study found that *Cinnamomum zeylanicum* and *Rosmarinus officinalis*, when combined, had a synergic effect against fungi developing on pears, meaning that the antifungal effect of the EO mix was greater than the sum of the antifungal activities measured separately [24]. As a future venue, it would be interesting to test the potential synergic effect of *Cinnamomum cassia* and *Origanum compactum*.

### A. oryzae, *a good model to study aspergillosis*

In this study, we used *A. oryzae* as an in *vitro* model of aspergillosis. While other *Aspergillus* fungi such as *A. fumigatus* are usually preferred to model aspergillosis [1–3,5,6,8], *A. oryzae* presents numerous advantages.

First, many species of the genus Aspergillus may cause aspergillosis. This includes *A. fumigatus*, the main cause of aspergillosis, but also *A. oryzae* [5,6]. Furthermore, *A. oryzae* and *A. fumigatus* are closely related both in terms of genome [8] and protein expression [9] (**Table 1**).

Second, *A. fumigatus* is classified as a Class 2 pathogenic agent, which means that scientists who wish to manipulate it must work in research facilities of an equivalent biosafety level. On the contrary, *A. oryzae* is a Class 1 pathogenic agent: it requires minimal security facilities and equipment to be manipulated. Therefore, *A. oryzae* represents a good alternative *in vitro* model of aspergillosis when a biosafety level 2 facility is not available – for example: a low-income country, or a teaching environment.

Finally, many studies have shown the antifungal effect of our EO panel against other members of the genus *Aspergillus*, including *A. fumigatus* [14– 18,21,22], which suggests shared mechanisms of action across many *Aspergillus* species.

## Acknowledgement

Firstly, we thank Dauphine Boutte, Julie Calmet, Adrien Dardinier, Julie Donzier, Mathilde Ducoudray, Fanny Marceau, and Charlotte Visconti who worked on a preliminary version of this project. We also thank Valentina Gligorijevic, Estelle Mogensen, and all members of the Teaching Laboratory at Sup’Biotech for their technical help. Finally, we thank the Sup’Biotech Administration team who facilitated our access to the research facilities and equipment.

The authors declare no competing interest.

